# Proteolytic activity of surface exposed HtrA determines its expression level and is needed to survive acidic conditions in *Clostridioides difficile*

**DOI:** 10.1101/2024.03.08.584076

**Authors:** Jeroen Corver, Bart Claushuis, Tatiana M. Shamorkina, Arnoud H. de Ru, Merle M. van Leeuwen, Paul J. Hensbergen, Wiep Klaas Smits

## Abstract

To survive in the host, pathogenic bacteria need to be able to react to the unfavourable conditions that they encounter, like low pH, elevated temperatures, antimicrobial peptides and many more. These conditions may lead to unfolding of envelope proteins and this may be lethal. One of the mechanisms through which bacteria are able to survive these conditions is through the protease/foldase activity of the high temperature requirement A (HtrA) protein. The gut pathogen *Clostridioides difficile* encodes one HtrA homolog that is predicted to contain a membrane anchor and a single PDZ domain. The function of HtrA in *C. difficile* is hitherto unknown but previous work has shown that an insertional mutant of *htrA* displayed elevated toxin levels, less sporulation and decreased binding to target cells. Here, we show that HtrA is membrane associated and localized on the surface of *C. difficile* and characterize the requirements for proteolytic activity of recombinant soluble HtrA. In addition, we show that the level of HtrA in the bacteria heavily depends on its proteolytic activity. Finally, we show that proteolytic activity of HtrA is required for survival under acidic conditions.

## Introduction

When entering a host, pathogenic bacteria are exposed to a plethora of stress-inducing conditions. These may include low pH, lysozyme, antimicrobial peptides, osmotic stress, elevated temperatures, the complement system, exposure to immune cells and their products such as antibodies, and many others. These conditions often result in unfolding of proteins in the bacterial envelope, which induces membrane stress and can eventually lead to cell death [1]. The gut pathogen *Clostridioides difficile* deals in part with these stresses through its spore stage. The bacteria travel from one host to the next as spores, which are resistant to many of the aforementioned factors. However, once in the intestines, the spores germinate and vegetative cells need to survive the possibly lethal, local stress factors.

It is vital that bacteria remove unfolded proteins from the bacterial surface, as unfolding impairs function. Removal can be done either through proteolytic breakdown or refolding of the unfolded proteins. One of the proteins that has been described to both refold and degrade proteins in different bacteria is the protease/foldase HtrA (High temperature requirement A) [2, 3].

HtrA is ubiquitous in nature and consists of a trypsin-like serine protease domain and one or two PDZ domains (<u>p</u>ost synaptic density protein (PSD95), <u>D</u>rosophila disc large tumor suppressor (Dlg1), and <u>z</u>onula occludens-1 protein (zo-1). It can be present as an extracytoplasmic soluble protein, or it can be membrane bound, depending on its precise function in a particular organism (Figure 1). HtrA generally forms trimers that may arrange into higher oligomeric structures once activated by potential substrates, but monomers have been described in *Helicobacter pylori* [4]. The PDZ domains are involved in oligomerization, substrate recognition and/or regulation of protease activity [2].

**Figure 1:**
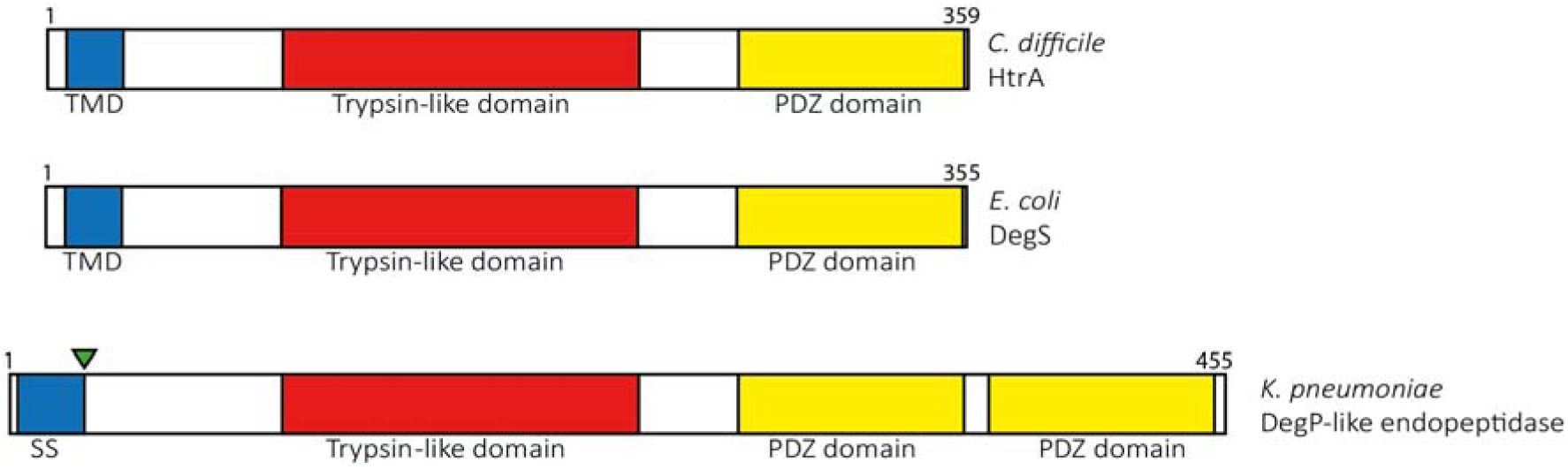
Schematic domain organization of *C. difficile* HtrA (CD3284), *E. coli* DegS and *K. pneumoniae* DegP-like endopeptidase. *C. difficile* HtrA and *E. coli* DegS both have one PDZ domain, and are anchored to the plasma membrane, whereas the *K. pneumoniae* DegP-like endopeptidase is a soluble, periplasmic protein. TMD: transmembrane domain; SS: signal sequence. Green triangle represents the signal peptidase cleavage site.

Previously, we have shown that an *htrA* insertion mutant in *C. difficile* demonstrates reduced spore formation, but a slightly higher virulence in a hamster model of acute infection. This is likely due to the increase of toxins produced by the mutant [5]. Furthermore, *in vitro* binding of *C. difficile* to Caco-2 cells was lower in the mutant. The increased virulence of the *htrA* mutant was unexpected, since other bacteria have been shown to become less virulent when they lack a functional HtrA. Deletion or inactivation of *htrA* in pathogenic bacteria often results in attenuation and loss of the capability to cause disease (virulence) [6–9]. Although it is clear that the lack of functional HtrA caused numerous changes in the *C. difficile htrA* insertion mutant, it is unclear whether the proteolytic activity or the chaperone/foldase activity underlie these responses.

In this manuscript, we investigate the localization and topology of the *C. difficile* HtrA protein. In addition, we study the requirements for proteolytic activity *in vitro* and examine the function of HtrA during stress. We show that 1) HtrA is extracytoplasmic and membrane bound , 2) *in vitro*, it displays proteolytic activity that is independent of the PDZ domain, 3) the proteolytic activity is important but the PDZ domain is dispensable for regulating HtrA expression levels in *C. difficile* and 4) the proteolytic activity and not the PDZ domain of HtrA is crucial for *C. difficile* to survive acidic conditions.

## Results

### Bioinformatic analysis predicts that HtrA consists of three functional domains

In bacteria, HtrA can be present in the gram-negative periplasm as soluble protease/foldase, like *Escherichia coli* DegP or DegQ or they can be membrane anchored, also extracytoplasmic, like *E. coli* DegS [10]. Some HtrAs are known to be secreted and are directly involved in virulence [11]. Bioinformatic analysis of *C. difficile* HtrA (CD3284, strain 630, uniport ID: Q180C8) predicts that it consists of a trypsin-like serine protease domain and one PDZ domain (http://smart.embl-heidelberg.de/). It is also predicted to a have an N-terminal hydrophobic domain (https://dtu.biolib.com/DeepTMHMM) that serves as a translocation signal, but it is not predicted to be cleaved after translocation (https://services.healthtech.dtu.dk/services/SignalP-6.0/), suggesting it to be an extracytoplasmic, membrane anchored protease, like *E. coli* DegS (see Figure 1) [12]. Alphafold (V2, Uniprot AF-Q180C8-F1-model_V4, average pLDDT score 85.70 over the whole protein, accessed February 2, 2024) [13] prediction and subsequent Foldseek analysis [14] shows the *Klebsiella pneumoniae* periplasmic DegP-like serine endoprotease (Uniprot A0A0H3GZ27) as the closest structural homolog (Sequence identity 33.7% and E-value 5.65 x 10^-33^; AFDB-Proteome). Remarkably, this is a soluble and periplasmic HtrA with two PDZ domains.

### HtrA is membrane-associated and localized in the extracytoplasmic space

In order to gain more knowledge about the working mechanism of HtrA in *C. difficile* and to confirm the bioinformatic analyses, we first decided to investigate its subcellular localization. Based on the above predictions, we hypothesized that HtrA would be present at the exterior side of the bacterial membrane, potentially located between the membrane and the cell wall, a space comparable to the periplasm in gram-negatives [15–18]. We applied differential centrifugation followed by anti-HtrA immunoblotting to study the localization of HtrA.

*C. difficile* WT cells (630Δ*erm*, JC049) were lysed using recombinant CD27L amidase (see experimental procedures) and subsequent sonication. The total lysate (Figure 2A, lane 1) was first centrifuged at 1300 x g to remove unbroken cells and cell debris (lane 2). The remaining part of the lysate (lane 3) was centrifuged at 200k x g to spin down membranes and peptidoglycan. The majority of the signal corresponding to HtrA was found in the pellet (Figure 2A, lane 5), and not the supernatant (lane 4) after centrifugation at 200k x g, but was solubilized when the pellet was treated with 2% Triton X-100 (lane 6). This suggests that HtrA is associated with membranes that are disintegrating upon treatment with detergent.

**Figure 2:**
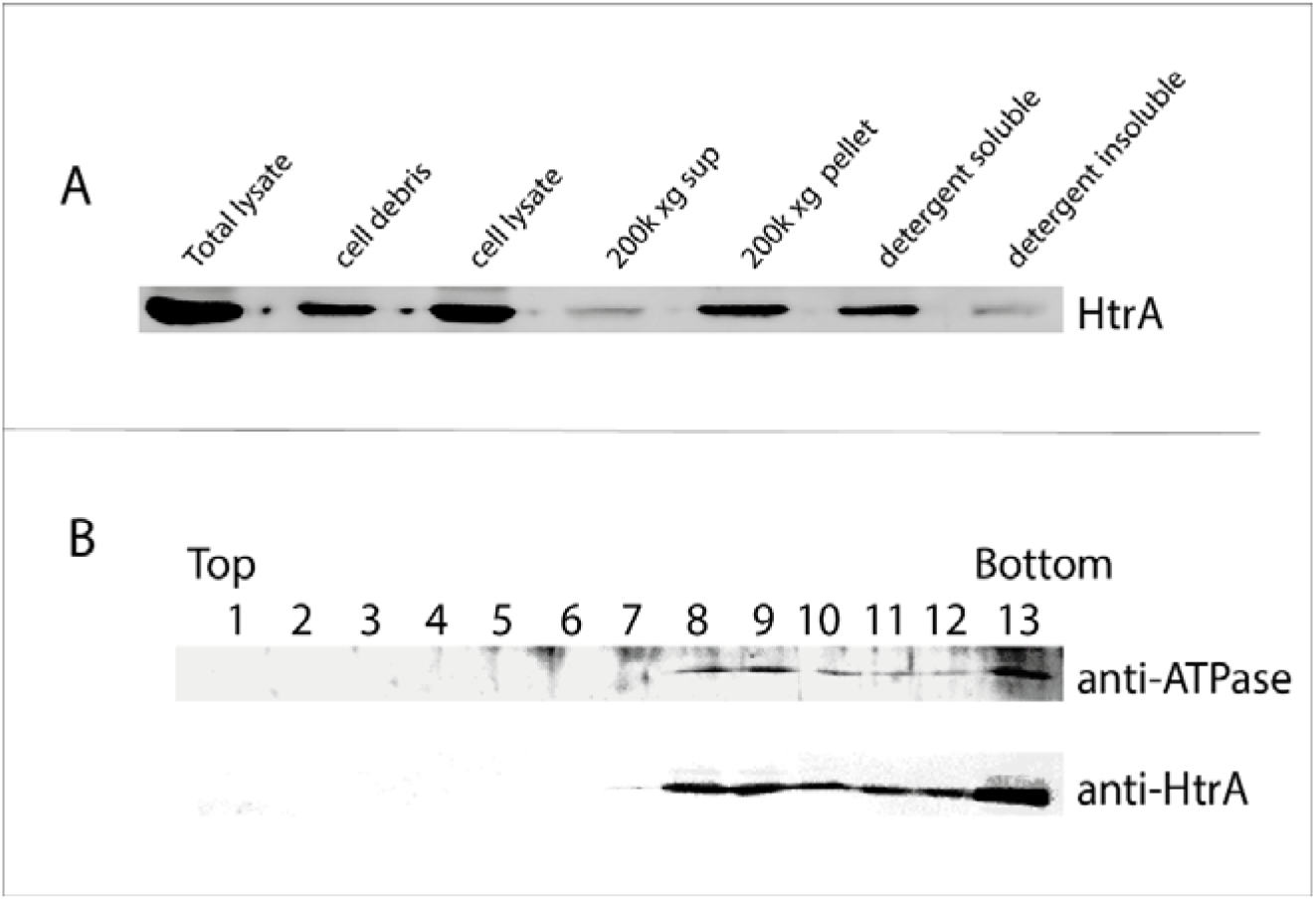
*C. difficile* HtrA is membrane-associated. (A) Immunoblot analysis of differential centrifugation of *C. difficile* lysates. Lanes (from left to right): Lane 1, Total lysate; Lane 2, cell debris, pelleted at 1300 x g centrifugation (unbroken cells); Lane 3, supernatant after 1300 x g centrifugation (cell lysate); Lane 4, supernatant after 200k x g centrifugation; lane 5, pellet after 200k x g centrifugation; The pellet was treated with detergent and centrifuged again at 20k x g. lane 6, detergent soluble proteins (supernatant after 20k x g centrifugation); Lane 7, detergent insoluble proteins (pellet after 20k x g centrifugation). (B) Immunoblot analysis of sucrose gradient fractions of fraction 5 of Figure 2A. ATPase F0F1 was used as a control for membrane proteins.

To further support this, the pelleted fraction after the 200k x g centrifugation step was loaded onto a 10-60% sucrose gradient. Following overnight centrifugation at 20k × g, fractions were collected and analysed by immunoblotting. HtrA co-sedimented with the integral membrane protein F0F1 ATPase [19] (Figure 2B). The presence of these proteins in the bottom fraction (fraction 13) is probably the result of aggregated material of the loaded sample (the pellet after the original 200k x g centrifugation). Altogether, the results in Figure 2 show that HtrA, in line with our prediction, is located in the bacterial membrane.

To determine the membrane topology of HtrA, we used the HiBiT Extracellular dDetection System (Promega, [20]). In this system, the NanoGlo LgBiT protein can be complemented with a HiBiT tag sequence, resulting in luciferase activity without lysis of the cells, provided the HiBiT tag is present exterior of the cell membrane. We cloned *htrA* with a C-terminal HiBiT tag, induced expression of this protein in *C. difficile* WT cells (JC049) and measured the luciferase activity. As controls, we included HiBiT-tagged HupA protein (CD3496), a cytoplasmic, nucleoid-associated protein [21] and HiBiT-tagged sortase (CD2718), a cell wall located protein [22, 23].

After induction, HtrA-HiBiT and sortase-HiBiT both showed a higher luciferase activity than HupA-HiBiT (Figure 3), indicating an extracellular localization of HtrA. The slightly elevated luciferase signals for the extracellular proteins at T0 are likely due to leakage of the inducible promoter, as was observed before [25]. A modest increase in luciferase activity was detected for HupA-HiBiT; we attribute this to cell lysis in combination with the highly sensitive detection by LgBiT. Luciferase activity in total cell lysates was comparable for all strains, indicating that the differences observed in Figure 3 are not the result of differences in expression levels (data not shown).

**Figure 3:**
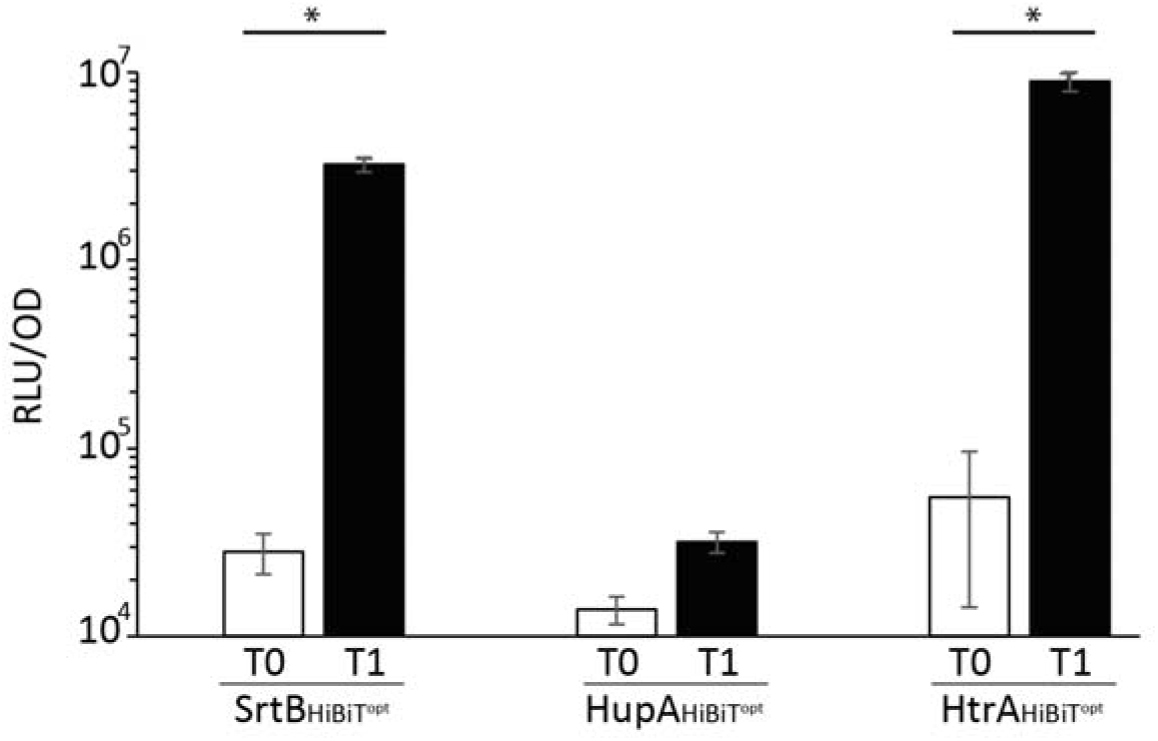
*C. difficile* HtrA is surface exposed. C-terminal HiBiT-tagged protein was expressed in *C. difficile* strain 630Δ*erm*, by inducing expression from P_tet_ with 200 nM anhydrotetracycline. Samples from the cultures were taken at the time of induction (white bars, T0) and 45 minutes after addition of anhydrotetracycline (black bars, T1) and used for measurement of luciferase activity using the HiBiT extracellular detection system. Luciferase values were corrected for the number of bacteria, as measured by OD_600nm_. The bars represent an average of four independent biological replicates with standard deviations. Data for SrtB and HupA were previously published [24] but stem from the same experiments as the HtrA experiments . * Significant difference T-test, P<0.001.

Together, the results in Figures 2 and 3 show that HtrA is located in the extracytoplasmic space, attached to the bacterial membrane, in line with the *in silico* prediction.

### HtrA protease activity is independent of the PDZ domain, but requires amino acids 30-55

In order to investigate the importance of the protease and PDZ domain for the proteolytic activity of HtrA, we thought to produce HtrA^S217A^ (lacking the serine in the catalytic triad) and HtrA lacking the PDZ domain. However, in order to obtain soluble HtrAs, we first of all cloned and recombinantly expressed HtrA lacking the N-terminal 64 amino acids (ΔN64), expected to contain the full protease and PDZ domain. Surprisingly, when HtrAΔN64 was tested using the HtrA model substrate β-casein [26, 27], no activity was observed. Assuming the transmembrane domain is not necessary for HtrA activity, this indicates that (part of) amino acids 30-64 are needed for proteolytic activity (Figure 4A). Investigation of the predicted HtrA structure and comparison with other HtrA structures showed that in the ΔN64 mutant, a small predicted α-helix between amino acid 56 and 64, was lacking. This small alpha helix (in green, Figure 4A) is conserved in most HtrAs, but is absent in the structure of trypsin. Hence, we tested a ΔN55 mutant, predicted to contain this α-helix, but this did not restore proteolytic activity, showing the importance of amino acids 31-54 (unstructured according to Alphafold V2 [13] predictions) for proteolytic activity of HtrA. As expected, deletion of only the transmembrane domain (ΔN30) yielded a proteolytically active form of HtrA which activity was abolished by mutating Ser-217 (Figure 4A) [5].

**Figure 4:**
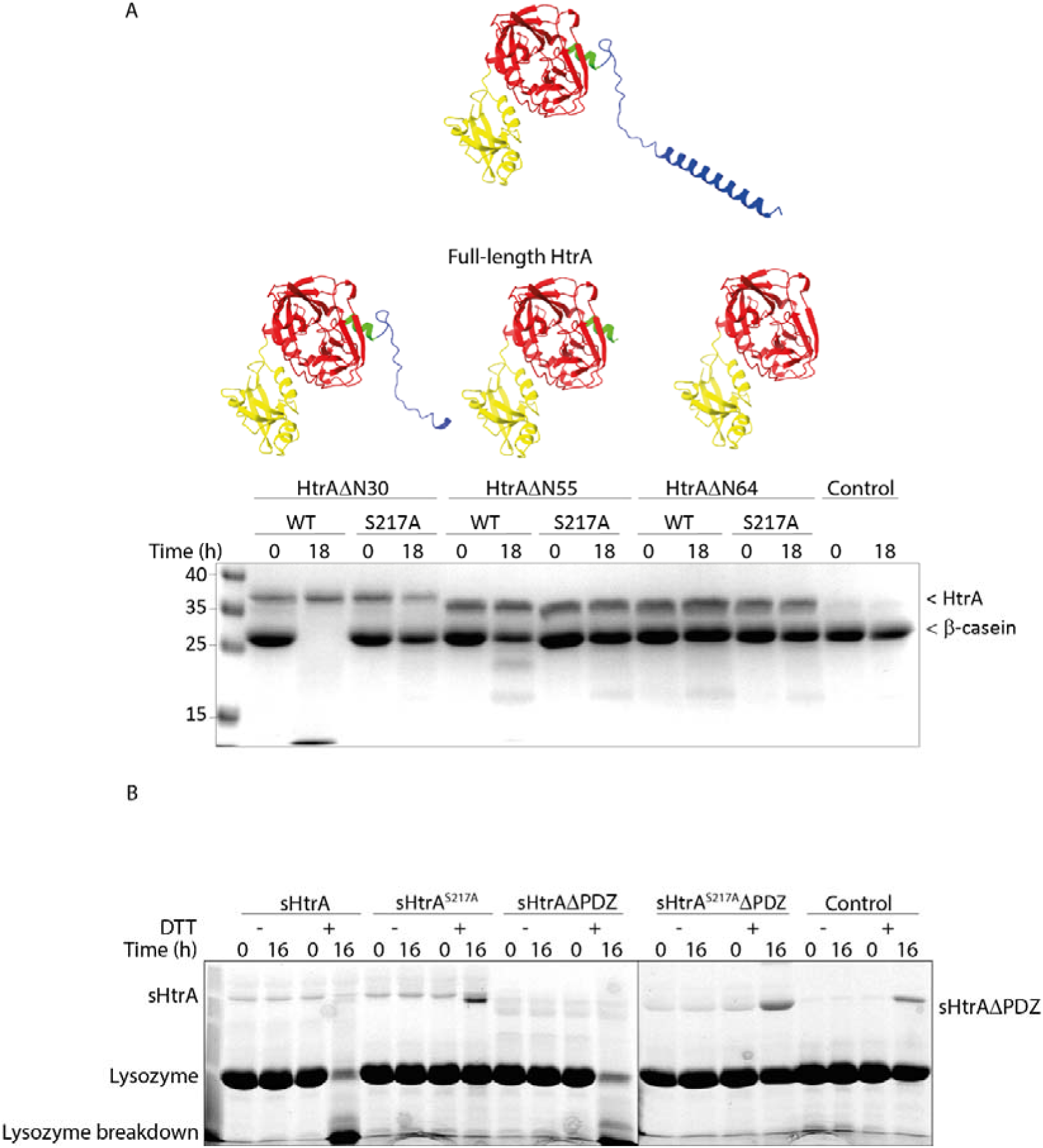
HtrA proteolytic activity is independent of the PDZ domain but requires the membrane-proximal amino acids 31-55. A) Several N-terminally truncated forms (ΔN30, ΔN55, ΔN64) of HtrA were incubated with β−casein analyzed on a 12 % SDS-PAGE, and stained with Coomassie Blue: N-terminus, including the transmembrane domain, Red: trypsin-like protease domain, yellow: PDZ domain, green: amino acids 56-64. HtrA models predicted by Alphafold V2. B) Lysozyme was incubated with various sHtrA forms in the presence or absence of DTT. Control lanes contain lysozyme without HtrA. Incubated mixes were analyzed on a Coomassie-stained 15% SDS-PAGE. The prominent band in DTT-treated, intact lysozyme lanes probably represents an aggregate or multimer of unfolded lysozyme. Samples at T=0 were taken after preparing the mixes, immediately mixed with loading buffer and stored at - 20 °C. All samples were taken during the same experiment, but run on two separate gels. The line represents the border between two separate gels.

Based on the results described above, we decided to use the HtrAΔN30 (sHtrA hereafter) and mutants thereof to further investigate properties of HtrA. Hence, we purified sHtrA, sHtrA^S217A^, sHtrAΔPDZ (ΔPDZ: deletion of amino acids 268-359) and sHtrA^S217A^ΔPDZ to >90% purity.

Since β-casein was approximately the same size as sHtrAΔPDZ, which would complicate interpretation of the gels, *in vitro* protease activity was measured using dithiothreitol (DTT)-treated (unfolded) lysozyme as a substrate. Unfolded lysozyme is used as an alternative model substrate to measure proteolytic activity of HtrA proteins [28] and has a predicted molecular weight distinct from all HtrA variants tested here (14 kDa). Since HtrA itself does not contain any cysteines, the folding of HtrA itself was not influenced by DTT.

As expected, lysozyme was degraded by sHtrA but not by sHtrA^S217A^ (Figure 4B) [5]. Interestingly, sHtrAΔPDZ also cleaved lysozyme, demonstrating the PDZ domain was not required for proteolytic activity *in vitro* (Figure 4B). This suggests that the PDZ domain is not needed to recognize potential substrates or that sHtrA does not require oligomerization through its PDZ domain to display proteolytic activity. Degradation of lysozyme solely happened in the presence of DTT, suggesting that sHtrA can only degrade unfolded protein.

### HtrA level is regulated by its own proteolytic activity in vivo

To further investigate the function of HtrA in *C. difficile*, we created a mutant that expressed a proteolytically inactive HtrA^S217A^ from its native promoter on the chromosome, using the Allele Coupled Exchange (ACE) method (Cartman et al., 2012). Surprisingly, this strain showed a dramatic increase of HtrA levels (Figure 5A). We sought to determine the source of the increased levels of HtrA in the *htrA*^S217A^ mutant through two lines for experiments. First, we determined the transcriptional activity of the P*htrA* promoter in various genetic backgrounds. Second, we determined protein levels in the *htrA*::CT mutant strain, complemented with different HtrA proteins expressed from the native *htrA* promoter. P*htrA* activity was monitored using a secreted, *C. difficile* optimized luciferase (sLuc^opt^) as a reporter molecule [30]. In the *htrA*^S217A^ strain and the *htrA*::CT strain, (Bakker et al., 2014) the luciferase activity was 37-fold and 17-fold higher, respectively, compared to WT (Figure 5B). This clearly shows that the increased HtrA level in the *htrA*^S217A^ strain (Figure 5A) is at least partially due to increased transcription and suggests that for maintaining a WT level HtrA, a feedback loop exists that involves the proteolytic activity of HtrA.

**Figure 5:**
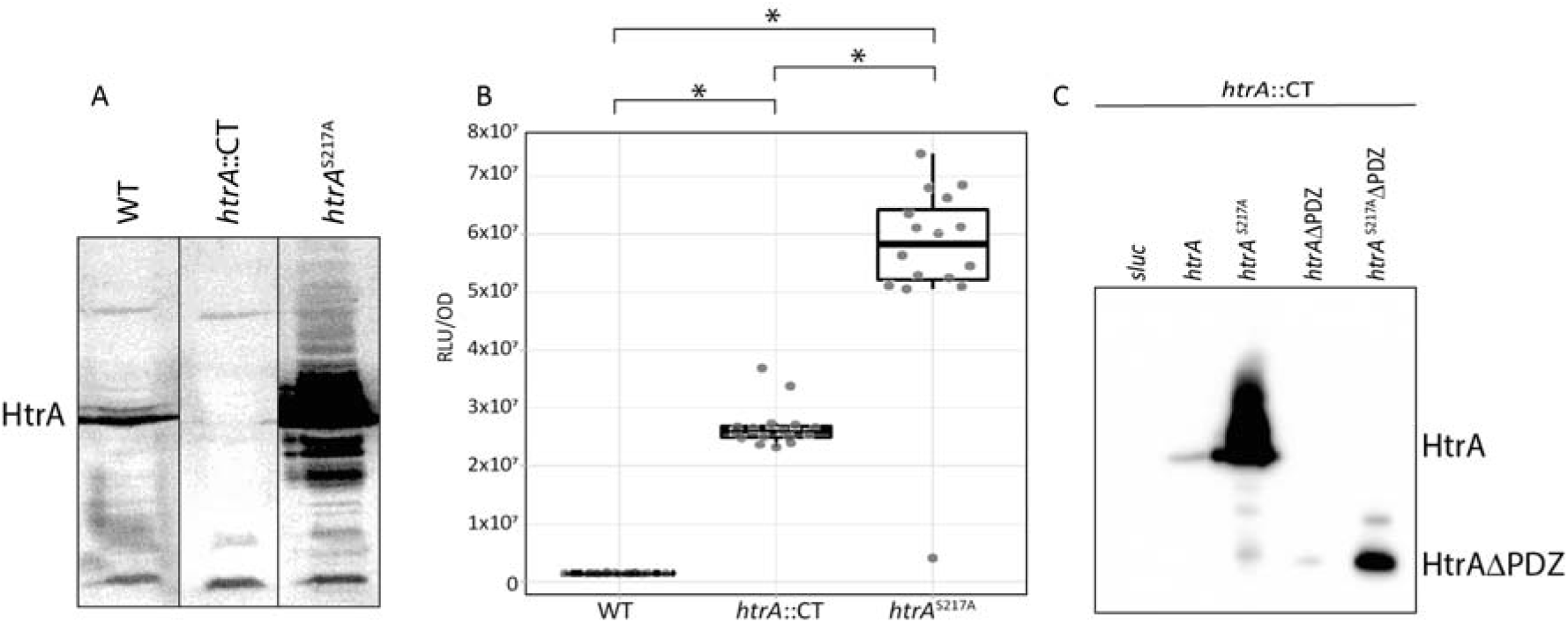
HtrA level is regulated in relation to its own proteolytic activity and is independent of its PDZ domain. A) The amount of HtrA in *C. difficile* cells was analyzed by immunoblot, using rabbit anti-HtrA serum. Lysates of equal amounts of cells were loaded on the gel, as determined by OD_600_ readings. B) transcription activity of P*htrA* was determined after three hours of growth in BHIY medium using a secreted form of NanoLuc (s*Luc*^opt^) as a reporter [30]. C) Complementation of the knockout strain with plasmids encoding P*htrA*-driven forms of *htrA* and a control *sLuc*^opt^, was carried out and amount of HtrA was determined by immunoblot. Equal amounts of cell lysates were loaded on the gel and immunoblots were probed with anti-HtrA serum.

In addition, we determined HtrA levels in the *htrA*::CT mutant, complemented by different forms of *htrA* to investigate which domain of HtrA is necessary to restore HtrA levels to wild type.

As expected, (Oliveira Paiva et al., 2016)complementation of the *htrA*::CT strain with WT *htrA* resulted in restoration of HtrA to WT levels (Figure 5C). In addition, complementation of *htrA*::CT with *htrA*Δ*PDZ* also resulted in WT level of HtrA (Figure 5C), demonstrating that the protease activity of HtrA is important for the complementation of this phenotype. In line with this, complementation of *htrA*::CT with *htrA*^S217A^ or *htrA*^S217A^ ΔPDZ did not result in WT HtrA levels, but showed persistent overexpression of HtrA, comparable to the *htrA*^S217A^ mutant (Figure 5A). This indicates that for maintenance of wild type levels of HtrA, the proteolytic activity of HtrA is required. Altogether, these experiments show that the absence of proteolytic activity, of HtrA, but not the PDZ domain, leads to up-regulation of *htrA* transcription.

### Comparative proteomics reveals differential expression of proteins in HtrA protease-deficient cells

To investigate possible differences in the levels of other proteins, we applied a comparative quantitative proteomic approach using light (*htrA*::CT), medium (WT) and heavy (*htrA*^S217A^) dimethyl labelling. Overall, 2210 *C. difficile* proteins with at least two tryptic peptides were found (Table S1). We decided to focus our further analyses on proteins that were found in all samples and showed consistent ratios. Hence, from the overall set of proteins, we selected only those proteins that, when comparing different strains, either had an abundance ratio above 2, below 0.5 or between 0.5 and 2 in all three biological replicates (Table S1). The resulting Volcano plots are presented in Figure 6.

**Figure 6:**
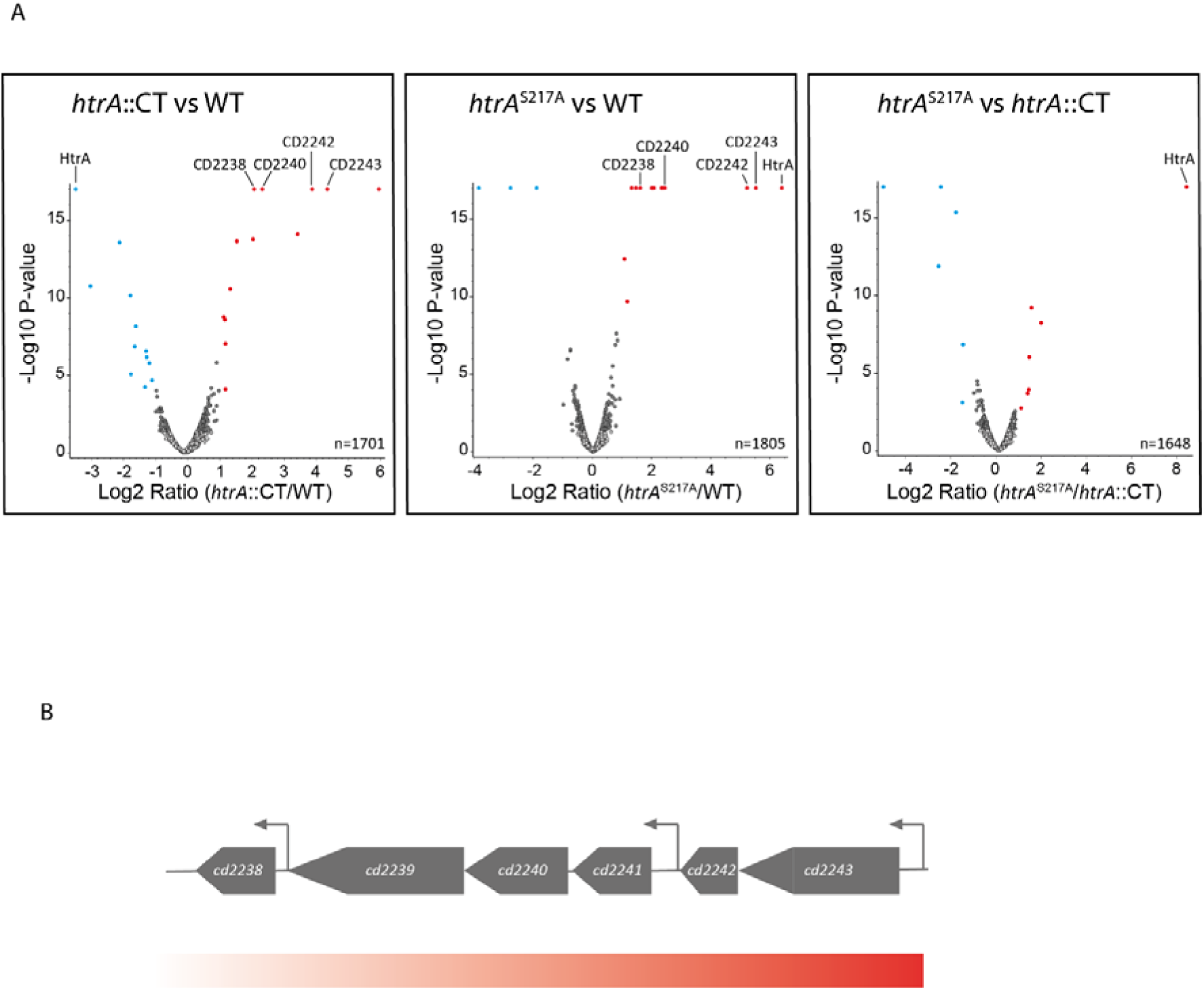
Comparative proteomics reveals upregulation of a specific set of proteins. A) Relative levels of proteins were determined through comparative proteomics. Ratio of proteins in indicated strains were visualized in volcano plots. Volcano plots of *C. difficile* strains show upregulation of a particular set of proteins in absence of HtrA proteolytic activity Dots in red indicate proteins with ratios >2 and dots in blue represent proteins with ratios <0.5. Ratios are an average of three independent experiments. B) Genomic region of genes coding for upregulated proteins. Arrowheads indicate tested promoters (P*cd2243*, P*cd2241*, P*cd2238*), gradient indicates level of upregulation, based on proteomics data, red is high, white is low.

First of all, HtrA was readily identified (18 unique peptides) and our quantitative proteomics data confirmed the knockout phenotype, even though some signal was found in the *htrA*::CT cells, especially in the data from one of the replicates). Importantly, the proteomics data also confirmed the high upregulation of HtrA in the *htr A*^S217A^ mutant cells shown above (Figure 5A). Based on the proteomics data, the levels of HtrA in the *htrA*^S217A^ cells was approximately 80-90 higher than in WT cells. Concurrently, even higher HtrA ratios were observed when comparing *htrA*^S217A^ and *htrA*::CT cells.

Interestingly, in both the *htrA*::CT mutant and *htrA*^S217A^ mutant cells, several other proteins were also highly upregulated. The highest upregulation was found for CD2243 (4 unique peptides, Uniprot ID: Q185B7) and CD2242 (9 unique peptides, Uniprot ID: Q185B8). Both proteins are encoded in a single operon. In addition, several other proteins (e.g. CD2240 (nanA), 13 unique peptides, Uniprot ID: Q185B3 and CD2238, 12 unique peptides, Uniprot ID: Q185A8) were found to be more abundant in the *htrA*::CT and *htrA*^S217A^ mutant cells, as compared to WT cells. These proteins are encoded in the same genomic region as the CD2242-CD2243 operon. We also note that for several other proteins encoded in the same genomic region indications for upregulation were observed (Table S1) but due to our stringent filtering routine described above, these were not considered in our data analysis.

The genes encoding the upregulated proteins CD2238-CD2243 (among which are the putative NanA and NanE) are not predicted to form a single operon. We tested the activity of three putative promoters, P*cd2243*, P*cd2241* (upstream of *nanE*) and P*cd2238*, in the wild type, *htrA*::CT and *htrA*^S217A^ strains to further investigate this. The data showed that only P*cd2243* was upregulated in the absence of HtrA proteolytic activity (Figure 7, Figure S1). P*cd2243* showed the highest activity in the *htrA*^S217A^ mutant, followed by the knockout and the wildtype, as was also observed for P*htrA* (Figure 5B). The lack of altered P*cd2241* and P*cd2238* activity (Figure S1) suggests that the elevated levels of proteins NanE, NanA and CD2238 in the *htrA*::CT strain and the *htrA*^S217A^ strain may be the result of transcriptional readthrough, likely caused by the high activity of P*cd2243* in these strains.

**Figure 7:**
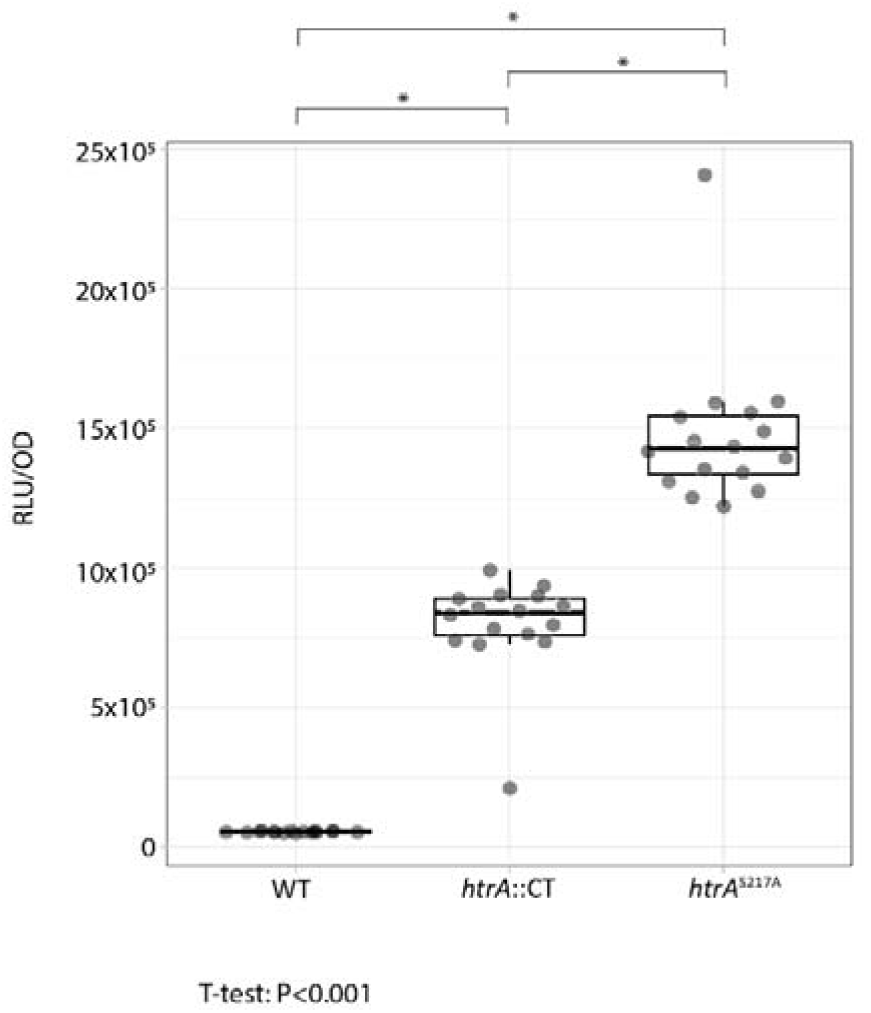
P*cd2243* shows differential expression in strains lacking HtrA proteolytic activity, similar to P*htrA*. Transcription activity of P*cd2243* was determined after three hours of growth in BHIY medium using a secreted form of NanoLuc as a reporter [30]. * Significant difference, P<0.001, T-test.

### Proteolytic activity of HtrA is required to resist low pH

In other bacteria HtrA is needed to survive exposure to stress conditions [6]. Therefore, we exposed wild type *C. difficile* to several stress conditions (low pH, H_2_O_2_, ethanol, high salt concentration) and measured the activity of P*htrA* under these conditions. Exposure to low pH induced increased P*htrA* activity, compared to the no stress control. All other applied stresses did not appear to induce elevated P*htrA* activity (Figure 8). Activity of P*cd2243* was increased similarly after exposure to low pH (Figure S2), once again suggesting that the activity of P*htrA* and P*cd2243* is regulated through a shared mechanism.

**Figure 8:**
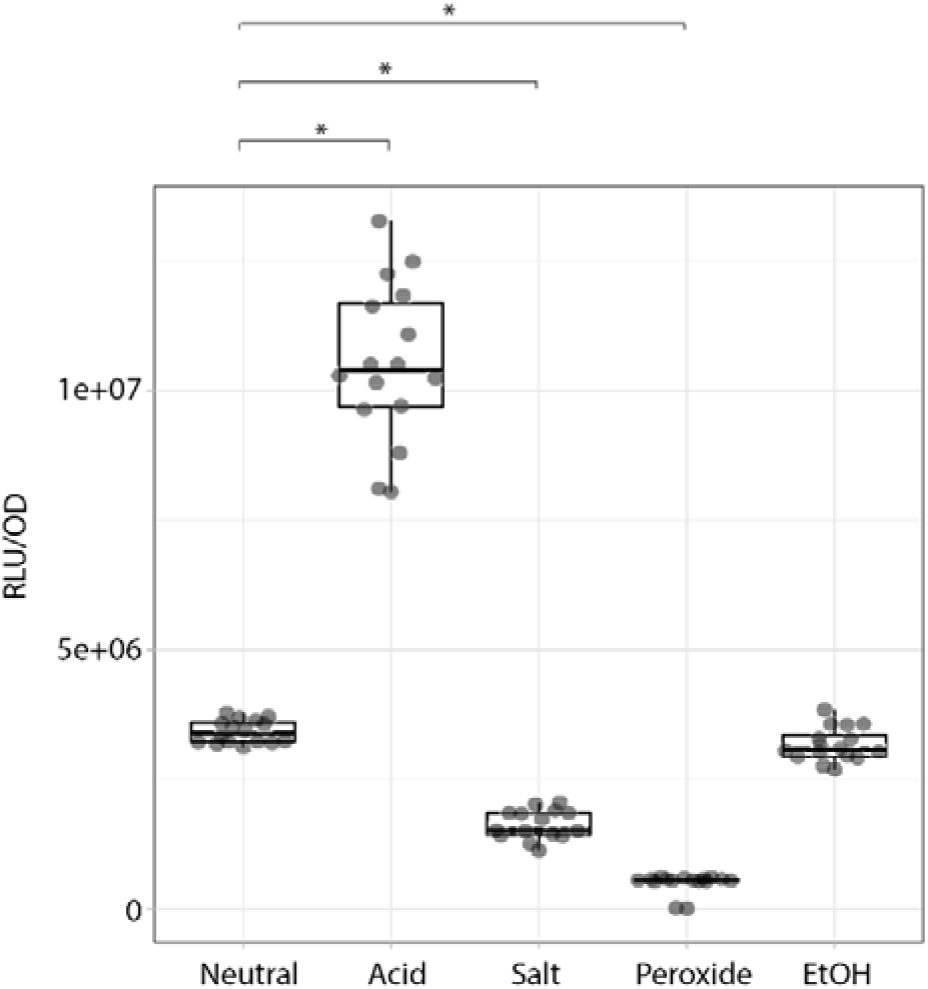
P*htrA* activity is increased during exposure to low pH. *C. difficile* WT, containing a P*htrA*-sLuc^opt^-containing plasmid was grown in BHIY + 15 µg/mL Thiamphenicol to an OD_600_ of approximately 0,3. Then, 200 µL of the culture was transferred to a 96 well plate and a stress was applied: pH 4.5 (Acid), 5% NaCl (Salt), 0.625% hydrogen peroxide (Peroxide), 3.2% ethanol (EtOH), or control (Neutral). Luciferase activity (Luc) was measured 90 minutes for 16 independent biological replicates after application of the stress and corrected for the optical density (OD) of the culture. The summary of the data is shown as a boxplot, with the box indicating the interquartile range (IQR), the whiskers showing the range of values that are within 1.5*IQR and a bold horizontal line indicating the median *Significant difference, P<0.001, T-test.

The up-regulation of *htrA* transcription at low pH suggested that HtrA expression may be relevant to survive acid stress. To test this, we applied filter discs, soaked in 10 % HCl to plates that were inoculated with the strains that expressed various forms of HtrA. After overnight incubation at 37 °C under anaerobic conditions, we measured the size of the halos around the filter discs. All strains that did not express proteolytically active HtrA, *i.e.* JC048 (*htrA*::CT), JC143 (*htrA*^S217A^), JC148 (*htrA*::CT, complemented with *htrA*^S217A^) and JC289 *(htrA*::CT, complemented with *htrA*^S217A^ ΔPDZ) showed larger halo sizes than the strains that did express proteolytically active HtrA (Figure 9A), irrespective of the presence of the PDZ domain. This is also evident when data from these strains are considered together; strains encoding proteolytically active HtrA showed a significant smaller halo size compared to strains lacking HtrA or expressing only HtrA^S217A^ (Figure 9B). In contrast, when a comparison was made between pooled strains encoding HtrA with or without the PDZ domain, no difference was found (Figure 9C). Thus, for better survival of acidic conditions, proteolytic activity of HtrA is essential and the PDZ domain is not.

**Figure 9:**
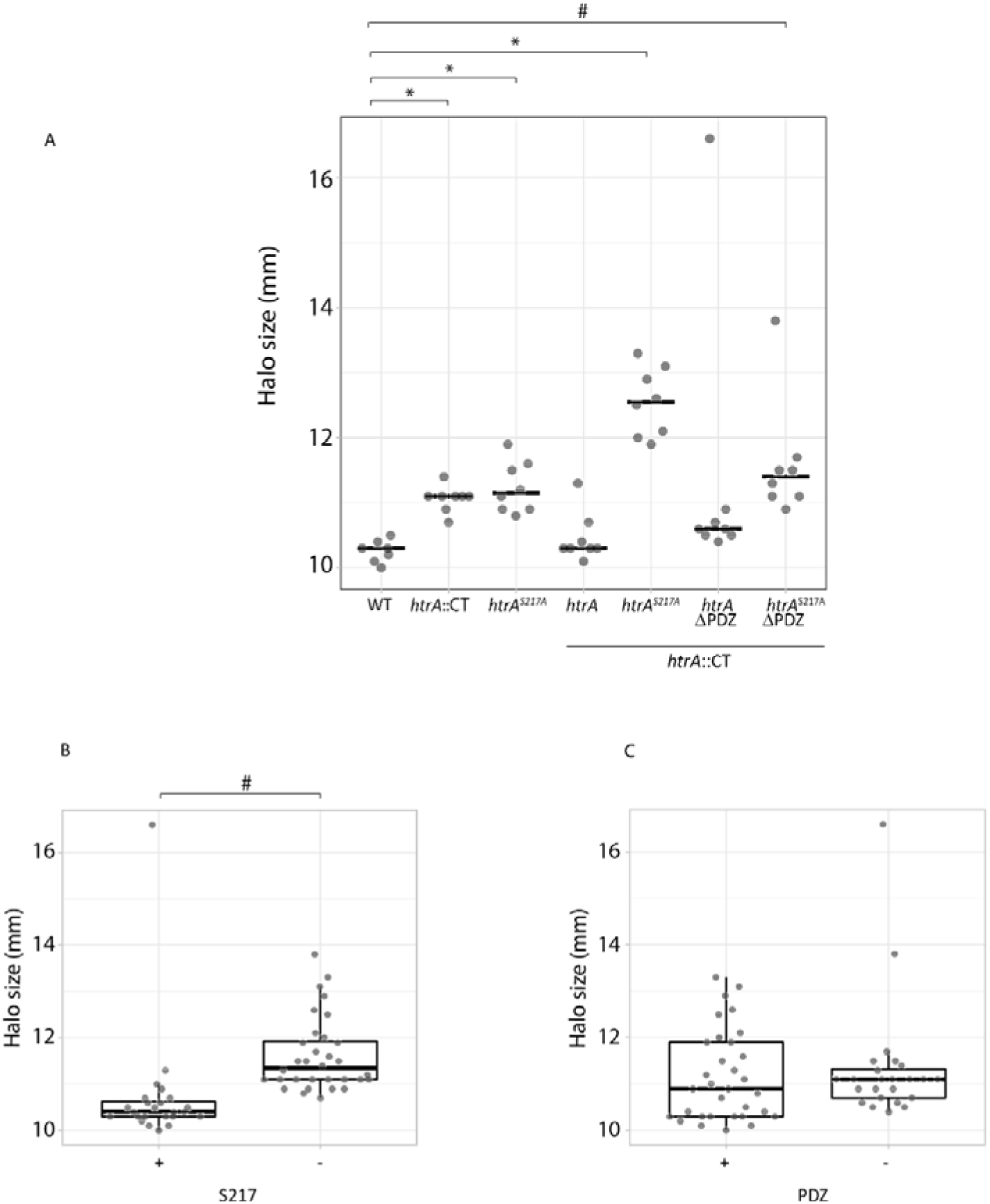
Proteolytic activity of HtrA is needed to resist low pH. A disc diffusion test with discs containing 10% HCl was carried out with strains expressing various forms of HtrA, controlled by P*htrA*. After 24 hours of growth, halos around the discs were measured. A) Sizes of halos for each strain are shown, with a horizontal line indicating the median of 8 biological replicates. B) Comparison between strains with a proteolytically active and inactive HtrA. C)Comparison between strains with HtrA with and without the PDZ domain. The summary of the data is shown as a boxplot, with the box indicating the interquartile range (IQR), the whiskers showing the range of values that are within 1.5*IQR and a bold horizontal line indicating the median.*significant difference, P<0.001; ^#^significant difference, P≤0,005, T-test.

## Discussion

In bacteria, homologues of HtrA exist in several forms with various subcellular locations. Differences are found in the presence/absence of a membrane anchor and the number of PDZ domains [2]. Based on analysis of the primary sequence, *C. difficile* HtrA is predicted to have a membrane anchor and one PDZ domain, similar to *E. coli* DegS [2]. Indeed, we show that *C. difficile* HtrA is membrane bound but present extracellularly (Figures 2 and 3). HtrA in *Bacillus anthracis* has a comparable domain organization, but has also been shown to cleave itself, ending up as a secreted protein [31]. We believe, however, that *C. difficile* HtrA is not secreted, since it was not found during a previous proteomics analysis of culture supernatant [32].

At present, it is unknown what the native substrates of HtrA in *C. difficile* are. The localization of HtrA limits the number of possible substrates relative to an intracellular localization, since the number of surface-exposed or secreted proteins is lower than the number of cytosolic proteins. During the in vitro assays with sHtrA we observed proteolytic activity using non-native substrates, but the rate at which this occurred was substantially slower than other HtrA-like proteins, like DegP or DegQ [2, 10]. This, combined with the membrane localization, suggests that HtrA in *C. difficile* may not be a protease/foldase involved in general protein homeostasis, but rather may have a regulatory role and (a) specific –as of yet unidentified– substrate(s).

We found a dramatic upregulation of HtrA expression in the chromosomal *htrA*^S217A^ mutant, which encodes a proteolytically inactive variant of HtrA. This suggests that the lack of proteolytic activity is somehow sensed -maybe through a feedback-loop-, causing a remarkable increase of promoter activity (Figure 5B). To our knowledge, the strong effect of mutagenesis of the active site of HtrA on its level of expression has not been observed in other bacteria. P*htrA* upregulation was also noticed in the *htrA*::CT strain. However, the activity of P*htrA* in the *htrA*^S217A^ strain was higher than in the *htrA*::CT strain. This might be caused by the accumulation of HtrA^S217A^ protein in the bacterial envelope, resulting in envelope stress which even further increases activity of P*htrA*.

Restoration of P*htrA* activity (and concurrent HtrA protein levels) occured when the strains were complemented with proteolytically proficient HtrA. However, the PDZ domain of HtrA was not necessary for this complementation (Figure 5C). Indeed, the PDZ domain was not needed for *in vitro* proteolysis by recombinant HtrA (Figure 4B). Moreover, the PDZ domain did not appear to be important for the role of HtrA in acid resistance (Figure 9). Several functions have been described for the PDZ domain(s) of HtrA [2, 33, 34]. In its inactive state, the membrane-bound *E. coli* DegS proteolytic activity is inhibited by the PDZ domain. Once unfolded outer membrane proteins bind to the PDZ domain, conformational changes trigger the proteolytic activity of the protease domain [33–36]. Consequently, DegS without PDZ is constitutively active [35]. DegS has no chaperone/folding activity, in contrast to DegP and DegQ, two soluble HtrAs, present in the *E. coli* periplasm. DegP and DegQ each contain two PDZ domains that are involved in recognition of substrates (*i.e.* unfolded proteins), which induces remodeling from an inactive hexamer to an active 12 or 24-mer [28, 37]. In *B. anthracis*, the HtrA PDZ domain is required for NprA secretion, but not for general stress resilience or even virulence in a murine model [31]. The HtrA PDZ domain in *C. difficile* might be involved in recognition of the native substrate(s) of HtrA and possibly in inhibition of HtrA proteolytic activity under non-stress conditions, a role similar to that described for *E. coli* DegS. In addition, DegS and *C. difficile* HtrA share the architecture of the protein with a membrane anchor and one PDZ domain, which could be indicative for a similar role in the bacterial life-cycle. With respect to the function of HtrA in the *C. difficile* life-cycle, it appears to play a role in withstanding acid exposure, like has been shown for HtrAs in other bacteria [6], but growth under mildly acidic or osmotic stress was not influenced (D. Bakker, W.K. Smits and J. Corver, unpublished results).

We noticed that truncating the N-terminus more than just the membrane anchor yielded a proteolytically inactive HtrA (Figure 4B). Alphafold [13] does not predict a specific structure for the amino acids 30-55, but they seem indispensable for proper proteolytic activity. This is reminiscent of the situation in *Helicobacter pylori* [38] where HtrA needs an amino acid stretch proximal to the N-terminus to form proteolytically active trimers. So far, we have been unable to show that HtrA from *C. difficile* forms trimers, using size exclusion chromatography of purified HtrA-ΔN30, which lacks the membrane anchor (results not shown). It is possible that the transmembrane anchor of *C. difficile* HtrA is involved in trimerization, or it does not trimerize at all.

It is interesting to note the increase in both P*htrA* and P*cd2243* activity during exposure to low pH and in the absence of proteolytic activity of HtrA, which suggests a functional link between HtrA and CD2242-CD2243. However, the increase in activity of both promoters differed greatly between the two conditions. Low pH induced a 1.5-2 fold increase in promoter activity, whereas absence of proteolytic activity resulted in 17-37 (P*htrA*) and 14-26 fold (P*cd2243*) increase. Interestingly, in vitro protease activity of sHtrA appears to be increased at pH 5.5 [5]. Perhaps then, the difference between the two conditions is explained in part by the increased activity of HtrA under acidic conditions, which cannot occur in the HtrAS^217A^.

Data from comparative proteomics (Figure 6) and reporter assays (Figure 5 and 7) suggest that *htrA* and the *cd2243* transcriptional unit are co-regulated but we did not identify similarities in the promoter sequences of *htrA* and *cd2243* that could explain their co-regulation. The function of CD2242 and CD2243 is unknown, but both are predicted to have a zinc-ribbon domain. In addition, CD2243 is predicted to have eight transmembrane helices (https://dtu.biolib.com/DeepTMHMM) with the zinc ribbon located in the N-terminal cytoplasmic domain (aa 1-92; https://phobius.sbc.su.se/), whereas CD2242 is predicted to be a soluble cytoplasmic protein.

We have previously compared transcriptional activity of the WT and the *htrA*::CT strains, using microarray analysis and did not find upregulation of *cd2243* or *cd2242*. Of the adjacent genes only *nanE* was upregulated approximately two-fold in both log phase and in stationary phase [5]. However, reanalysis of that raw data shows that indeed both *cd2242* and *cd2243* were highly upregulated during logarithmic phase, but not at stationary phase. Due to stringent filtering (related to high variance), these data did not end up in the previous manuscript. Since our proteomics data and the data obtained with the reporter analysis both were carried out at stationary phase, the results can be considered as a cumulative effect of the upregulation during the logarithmic phase.

In conclusion, we describe *C. difficile* HtrA as a membrane bound extracellular protein that displays proteolytic activity. This activity is important for regulation of HtrA expression levels and for survival under acidic conditions, suggesting a role for HtrA in survival of *C. difficile* in the host, as shown for HtrAs in other bacteria.

Previously, we have shown that the *htrA*::CT strain was more virulent in a hamster model of *C. difficile* infection [5], probably due to increased toxin production in this strain, combined with the exquisite sensitivity of the hamsters for *C. difficile* toxins. Other bacteria are known to display a less virulent phenotype in absence of HtrA, caused by the inability to survive stresses encountered in the host [6]. Testing the HtrA mutants in an animal model that is less sensitive for the toxins might reveal a similar role for HtrA in *C. difficile*.

## Experimental procedures

### Bacterial culture and media

*E. coli* strains were cultured in LB broth (Sigma), using the appropriate antibiotics.

*C. difficile* cultures were grown on plates consisting of Brain heart infusion (Oxoid), supplemented with 0,5% yeast extract (Sigma) (BHIY) and 1,5% agar (Thermo). Antibiotics were added when needed. Liquid cultures were grown in BHIY, supplemented with antibiotics when necessary.

All strains used in this study are listed in Table S2.

### Anti HtrA serum

Recombinant HtrA was made in *E. coli* as previously described [5]. 200 µg was injected into a rabbit and after three boosters (in 3 months), the serum was yielded (Eurogentec).

### Membrane association

Membrane association was assessed essentially as [19], with a few modifications. Five-mL aliquots of overnight cultures of *C. difficile* were harvested by centrifugation at 4000 g. Pellets were washed with PBS and subsequently resuspended in 500 µL 50 mM Tris, pH 7,4 + cOMPLETE protease inhibitor cocktail EDTA-free (1 tablet/50 mL) (Roche). Recombinant CD27L amidase was added to the bacterial suspension at 30 µg/mL, followed by 2hr incubaton at 37 °C. The expression plasmid encoding the amidase (pHAS042) was a kind gift of Dr. Robert Fagan [39]. Thereafter, the bacterial suspension was sonicated 3x for 30 s at 10 microns amplitude. Subsequently, DNAse (100 µg /mL) and RNAse (100 µg /mL) were added and the samples were incubated on ice for 30 minutes, after which a 5 µL sample (sample 1) was taken and stored. The rest of the suspension was centrifuged for 5 minutes at 1300 × g, to remove unbroken cells. The pellet was resuspend in 50 µL 50 mM Tris, pH 7.4 and a 2 µL sample was taken and stored (sample 2). Of the remaining supernatant, a 5 µL sample was taken and stored (sample 3). Subsequently the supernatant was split in two parts and centrifuged at 200k × g for 60 minutes at 4 °C. A 5 µL sample was taken from the supernatant and stored (sample 4). The two pellets were processed separately. 1) The pellet was resuspended in 200 µL 50 mM Tris, pH 7.4, 5 mM EDTA and 2% triton X-100, after which it was incubated for 30 minutes at room temperature. A 2 µL sample was taken and stored for analysis (sample 5), after which the remaining solution was centrifuged for 1 hr at 20k x g at 4 °C. A sample of 2 µL was taken from the supernatant (sample 6) and stored for analysis. The pellet was resuspended in 50 µL Tris buffer and a 5 µL sample was taken and stored for analysis (sample 7). All samples were analyzed for HtrA content by immunoblot, using a rabbit anti-HtrA serum. 2) The other ultracentrifuge pellet was resuspended in 200 µL PBS and loaded on a sucrose gradient (TLS 55 centrifuge tube) which consisted of 200 µL layers of 60, 50, 40, 30, 15 and 10% sucrose (in PBS). The gradient was spun at 200k × g for approximately 18 hours and subsequently fractionated in 200 µL fractions. Twelve µL of each fraction was analyzed by immunoblot, using a rabbit anti-HtrA serum.

### Luciferase reporter assay

Hundred µL of a 100-fold diluted culture of *C. difficile* strains carrying sLuc^opt^ reporter constructs was mixed with 20 µL of reconstituted Nanoglo substrate (50 fold diluted Nanoglo substrate in kit buffer, Promega) in a white, flat bottom 96-well plate. Subsequently, relative light units (RLU) were measured in a GloMax® Explorer Multimode Microplate Reader (Promega), using standard settings. RLU was corrected for biomass, by dividing the RLUs through the OD_600nm_ value of the bacterial culture at the time of harvest.

### HiBiT extracellular detection system

The assay was carried out essentially as described [24]. In brief, 50 µL of cultured *C. difficile* was mixed with 50 µL kit reagents (100-fold diluted LgBiT protein, 50-fold diluted substrate in kit buffer, Promega). Subsequently, the reactions were incubated for 10 minutes at room temperature, after which RLUs were measured in a white flat-bottomed 96-well plate, using a GloMax® Explorer Multimode Microplate Reader (Promega).

### Conjugations

Conjugations were carried out as previously described [40]. Briefly, *E. coli* strain CA434 was transformed with the desired plasmid and selected for on LB plates containing 20 µg /mL chloramphenicol. A single colony was grown overnight in LB/20 µg /mL chloramphenicol. A pellet of 1 mL of culture from the *E. coli* CA434 transformants was imported in the anaerobic chamber (VA-1000 or A55 HEPA workstation, Don Whitley Scientific) and mixed with 200 µL of an overnight culture of *C. difficile*. This mixture was drop-wise plated on a BHI/0.5% yeast plate and incubated for >6 hrs at 37 C under anaerobic conditions. Subsequently, the bacteria were scraped of the plate into anaerobic PBS and dilutions were plated on BHIY plates, containing 15 ug/mL thiamphenicol (BHIY/Thia) and *C. difficile* selective supplement (CDSS, Oxoid). Colonies growing were passaged three times on the same plates and were eventually tested for the species and presence of the plasmid by PCR, using primers oWKS1070/oWKS1071 (specific for *C. difficile gluD*) and oWKS1387 and oWKS1388 (specific for *traJ*, present on the plasmids) .

### Cloning

All primers used for cloning are listed in Table S3. We used a PCR-based approach to generate expression constructs for HtrA-variant proteins based on published plasmids pJC014 and pJC051 [5], encoding *C. difficile* sHtrA and sHtrA^S217A^. To enable recombinant expression of sHtrAΔPDZ and sHtrA^S217A^ΔPDZ we generated pJC105 and pJC109 using primers CD3284F2 and CD3284R2 and pJC014 and pJC051 as templates for PCR. Likewise, we used primers CD3284F5 and CD3284R3 to clone pJC057 and pJC058 and primers CD3284F6 and CD3284R3 were used to clone pBC013 and pBC015.

To generate a P*htrA*-*sluc*^opt^ reporter construct, we used primers CD-PHTRAF2 and CD-PHTRAR2 to amplify the promoter of *htrA*. Subsequently, it was cloned into pAP24 [30] using restriction enzymes KpnI and SacI, yielding pJC085. Plasmids pJC086, pJC087 and pJC088 were similarly constructed using primers CDPCD2243F and CDPCD2243R, CDPCD2241F and CDPCD2241R and CDPCD2238F and CDPCD2238R, respectively.

To construct P*htrA*-driven complementation plasmids, we used primers CD-PHTRAF2 and oDB0068 to amplify the WT *htrA* sequence including the *htrA* promoter. The amplicon was cut with KpnI and BamHI to clone it into pRPF185 [40], yielding pJC080. We used primers CD3284-S217AF and CD3284-S217AR to create pJC081 through QuikChange mutagenesis. pJC102 was constructed as follows: primers CD-PHTRAF2 and CD3284R2 were used to amplify the P*htrA-htrA*ΔPDZ. This amplicon was cloned into pCR2.1-TOPO (Thermo). Subsequently, this plasmid was cut with KpnI and BamHI and ligated into pRPF185. Primers CD3284-S217AF and CD3284-S217AR were used to create pJC115 through QuikChange mutagenesis

To create pJC113 (encoding HtrA-HiBiT), we used oDB0067 and CD3284R6 to amplify the *htrA* coding sequence. The amplicon was cut with SacI and XhoI and ligated into pAF302 [24] that was cut with the same enzymes.

Plasmid pJC076 was created using Gibson assembly using 3 fragments: fragment 1 was the pMTL-sc7315 backbone [29], amplified with primers oWKS1537 and oWKS1538, fragment 2 was the N-terminal part of cd3284 plus 1000 bp upstream of the S217 in *cd3284*, amplified with primers CD3284_N-TERMINUS_FWD and CD3284_N-TERMINUS_REV and fragment 3 was the C-terminal part of cd3284 plus 1000 bp downstream of the S217 in *cd3284*, amplified with primers CD3283_C-terminus_FWD and CD3284_C-terminus_REV. All three fragments were purified and combined in a Gibson assembly (New England Biolabs [41]). Sequencing revealed a point mutation in the cd3284 open reading frame, resulting in a T184I mutation. This mutation was restored back to wild type using primers CD3284ITF and CD3284ITR in a QuikChange reaction.

All PCRs needed for cloning were carried out with high fidelity enzymes Q5 polymerase (New England Biolabs) or Accuzyme (Bioline). All plasmids are listed in Table S4.

### Construction of chromosomal *htrA*^S217A^ mutant

pJC076 was transformed into *E. coli* CA434, which was then used as a donor in conjugation with 630Δ*erm*. Conjugation was carried out as described above. Transconjugants were passed three times onto BHIY/Thia plates supplemented with CDSS and subsequently analysed for single cross-over with primers oWKS1539 and CD3284DOWR. Positive clones were cultured on BHIY/CDSS plates to allow double cross for 48 hours. After that, cells were scraped of the plates with anaerobic PBS and dilutions were plated onto *C. difficile* minimal medium (CDMM) plates, containing 50 µg /mL 5-fluorocytosine (5FC Sigma). After 48 hr, colonies were plated on BHIY/Thia plates and CDMM-5FC plates. Colonies growing on CDMM-5FC plates, but not on BHIY/Thia plates were selected and DNA was isolated using a DNeasy blood and tissue kit (Qiagen) to investigate whether the intended chromosomal mutation was introduced. Eleven out of sixteen strains sequenced contained the desired mutation in *htrA*, resulting in an amino acid change serine to alanine of position 217 in the CD3284 protein (HtrA). One clone (JC143) was used for further analysis and experiments.

## Proteomics

### Filter-Aided Sample Preparation (FASP) and dimethyl labelling

*C. difficile* wild type (strain 630Δ*erm*), *htrA*::CT and the *htrA*^S217A^ strain (JC143) cells were harvested after 24 hours of culture (50 mL) at an OD600 of 0.6. Every pellet was resuspended in an equal volume of 0.2M TRIS-HCl pH 7.6 and then 4 volumes of ST lysis buffer (5% SDS, 0.1M TRIS-HCl pH 7.6) were added.

Cells were heat-inactivated by incubation at 95°C for 10 minutes and then mechanically lysed using a bead-beater (TissueLyser LT; Qiagen). Briefly, samples were incubated with glass beads (glass beads-acid washed; Sigma-Aldrich) for 5 minutes at a speed of 50 oscillations per second. After the first cycle of bead-beating, samples were put on ice for 5 minutes and the procedure was repeated twice. Eventually, the samples were spun down for 1 minute at 20000 x g, supernatants were collected and protein concentration was determined using the BCA assay (Thermo Fisher Scientific). An amount equivalent to 100 µg of proteins per sample was incubated with DTT at a final concentration of 0.1M for 4 minutes at 95⁰C and samples were further processed according to the FASP procedure previously described [42]. In brief, reduced proteins were loaded on a 30 kDa filter, washed with 8M urea to eliminate the SDS and alkylated with 50mM iodoacetamide. Then, proteins were digested overnight with Endoproteinase Lys-C (endoLysC, Sigma) using a total protein/enzyme ratio of 20: 1 (w/w) followed by a second 4-hour-digestion with trypsin (Worthington Enzymes Lakewood, NJ, United States), using the same total protein/enzyme ratio. For quantitative analysis, samples were labelled using stable isotope dimethyl labelling. Briefly, a C18-HLB 1 cc cartridge (Oasis) was first washed with 90% acetonitrile (ACN) and then equilibrated with 0.1% formic acid (FA) washing buffer. Subsequently, 50 µg of peptides from *C. difficile htrA*::CT cells were diluted in washing buffer to a final volume of 1 mL and applied to the cartridge. After washing, the “light labelling mixture” (4.5 mg sodium cyanoborohydride (NaCNBH_3_), 14 μl formaldehyde (CH_2_O) 37%/ 2.5 mL sodium phosphate buffer pH 7.5) was added during 5 minutes, applying 0.5 mL at 1 minute-intervals of incubation. Subsequently, the cartridge was washed again twice before the second sample (50 µg of peptides from *C. difficile* WT cells) were loaded and labelled with the “medium labelling mixture” (4.5 mg NaCNBH_3_, 25 µL deuterated formaldehyde (CD_2_O) 20% / 2.5 mL sodium phosphate buffer pH 7.5). The same procedure was then repeated with 50 µg peptides from *C. difficile htrA*^S217A^ cells using the “heavy labelling mixture (4.5 mg deuterated sodium cyanoborohydride, NaCNBD , 25 µL deuterated and ^13^C labelled formaldehyde (^13^CD O) 20% / 2.5 mL sodium phosphate buffer pH 7.5). Finally, after three washings with 0.1% FA, labelled peptides were eluted using 400 µL of an 80% ACN/ 0.1% FA solution. The whole procedure described above was performed with three biological replicates, resulting in three pooled samples of 150 µg labelled peptides.

### Strong cation exchange (SCX) chromatography

Before LC-MS/MS analysis, labelled samples were fractionated by SCX. In short, samples were resuspended in 20 µL solvent A (30% ACN, 0.1% FA) and fractionated by SCX using an Agilent 1100 system equipped with an in-house packed SCX-column (320 μm ID, 15 cm, PoIySULFOETHYL A 3 μm; Poly LC), at a constant flow rate of 3 μl/min. Peptide separation was obtained with a linear gradient from 0 to 100% of solvent B (70% 0.25M KCl, 30% ACN and 0.1% FA) over 15 minutes, followed by 15 minutes of 100% solvent C (70% 0.5M KCl, 30% ACN and 0.1% FA). 16 fractions were collected at 1 minute-intervals, freeze-dried and stored at -20⁰C until LC-MS/MS.

### LC-MS/MS

Freeze-dried SCX fractions were resuspended in 30 μl of a 95% water, 3% ACN, 0.1% FA solution and analysed by LC-MS/MS using an Easy nLC 1200 gradient HPLC system coupled with an Orbitrap Fusion Lumos Tribrid Mass Spectrometer (Thermo Fisher Scientific). Fractions (5 μL) were injected onto an in-house packed pre-column (100 μm×15 mm; Reprosil-Pur C18-AQ 3 μm; Dr. Maisch), equilibrated with solvent A (0.1% FA) and then eluted using a homemade analytical nano-HPLC column (15 cm×50 μm; Reprosil-Pur C18-AQ 3 μm), with a linear gradient from 10% to 40% solvent B (80% ACN, 0.1% FA) over 120 minutes, at a constant flow rate of 250 nl/min.

The Orbitrap Fusion Lumos Tribrid Mass Spectrometer was used in the data-dependent mode and MS spectra were collected from *m/z* 400 to 1500 (resolution 120.000, maximum injection time 50 ms). Higher-energy collisional dissociation (HCD) MS/MS at a normalised collision energy of 32% was performed (resolution 30.000, isolation window (*m/z*) of 1.2) on the 10 most abundant ion precursors (charge states 2-4), with a dynamic exclusion time of 1 min.

### Data analysis

For peptide identification, MS/MS spectra were searched against the UniProt *C. difficile* strain 630 database (3762 entries) using Mascot Version 2.2.07 (Matrix Science) with the following settings: 10 ppm and 20 milli mass units tolerance for precursor and fragment ions, respectively; trypsin was set as enzyme and two missed cleavages were allowed. Carbamidomethyl on cysteines was set as a fixed modification. Variable modifications were oxidation (on Met) and acetylation on the protein N-terminus. In addition, light (+28.0313), medium (+32.0564) and heavy (+36.0757) dimethyl modifications on Lys and the peptide N-terminus were set as variable modifications. All searches and subsequent data analysis, including Percolator and relative quantification, were performed using Proteome Discoverer 2.5 (Thermo Scientific). Peptide-spectrum matches were adjusted to a 1% FDR.

### Data availability

The mass spectrometry proteomics data have been deposited to the ProteomeXchange Consortium via the PRIDE [43] partner repository with the dataset identifier PXD050242.

## Recombinant protein expression and purification

Recombinant HtrA was expressed in *E. coli* and purified as previously described [5].

## Protease assays

To detect proteolytic activity, 0.25 nmol of HtrA was mixed with 2 nmol of substrate (β-casein or lysozyme) at pH 5.5 [5]. Immediately after mixing, a sample was taken (t=0) and the remaining mixture was incubated for 16 hours at 37 °C. Subsequently, a sample was taken and all samples were analyzed on a 12.5% (for β-caseine) or 15% (for lysozyme) SDS-PAGE. Gels were stained with Coomassie to visualize proteins.

## Filter disc assays

Two hundred µL of overnight *C. difficile* cultures in BHIY were plated onto BHIY plates, supplemented with appropriate antibiotics when necessary. Afterward, filter discs, soaked with 10 µL of 10% hydrochloric acid, were applied onto the plates. Subsequently, the plates were incubated for 24 hr at 37 °C in an anaerobic chamber (Don Whitley A55 HEPA) and the inhibition zones were measured with a Scan 500 colony counter (Interscience), using the inhibition zone mode.

## Data processing and preparation of figures

Numerical data was processed in Microsoft Excell and Plots of Data (https://huygens.science.uva.nl/PlotsOfData/). All figures were prepared in Adobe Illustrator.

## Acknowledgements

The authors wish to thank Sjaak van Voorden for help with cloning.

## Author contributions

J.C. designed and performed experiments, analysed and interpreted the data and wrote the paper, B.C. performed experiments and interpreted the data, T.M.S. performed experiments, interpreted the data and wrote the paper, A.H.R. performed experiments, M.L. performed experiments and interpreted the data, P.J.H. designed and performed the experiments, analysed and interpreted the data and wrote the paper, W.K.S. designed the experiments, interpreted the data and wrote the paper.

## Abbreviated summary

Absence of HtrA proteolytic activity leads to upregulation of *htrA* in *Clostridioides difficile* and a zinc-ribbon-containing membrane protein. HtrA is also upregulated during acid stress and the proteolytic activity is needed to survive these conditions. The HtrA PDZ domain is not needed for proteolytic activity or survival under acidic conditions.

