## Supplemental files figure S1, S2, table S2 for "Proteolytic activity of surface exposed HtrA determines its expression level and is needed to survive acidic conditions in *Clostridioides difficile*"

Corver et al.,

Supplemental figures S1 and S2. Tables S2, S3 and S4.

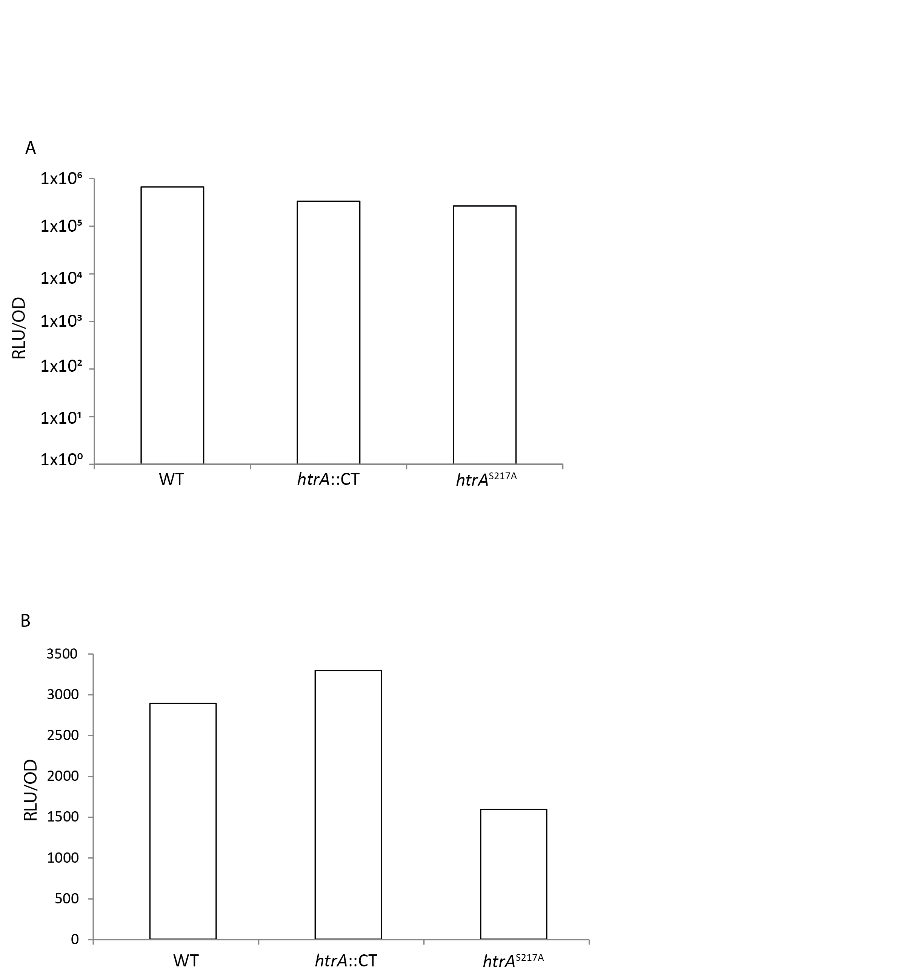

**Figure S1: Promoter activity of P*cd2241* or P*cd2238* is independent of HtrA activity.** Transcription was determined at early stationary phase after growing in BHIY. A) P*cd2241* activity, B)P*cd2238* activity. Bars are an average of two independent experiments.

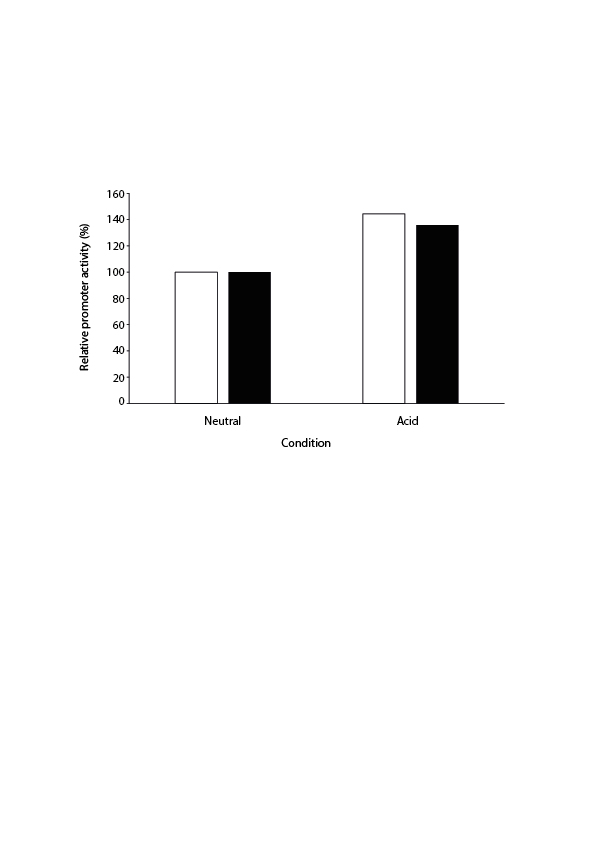

**Figure S2 : P*htrA* and Pc*d2243* are both upregulated after exposure of *C. difficile* to low pH.** In this experiment, both promoters were upregulated 1,4 x, compared to the neutral pH condition. Promoter activity was measured at neutral pH and after 90 minutes of exposure to low pH. The relative activity was calculated as a percentage of the activity at neutral pH. P*htrA* activity is shown in white bars, P*cd2243* activity is shown in black bars.

Table S2: *C. difficile* strains used in this study

| Strain | genotype | resistance | Reference |
| --- | --- | --- | --- |
| JC049 | 630Δ*erm* |  |  |
| JC048 | 630Δ*erm*, *htrA*::CT^a^ | *ermB* | DB002 [1] |
| JC143 | 630Δ*erm htrA^S217A^* |  | This paper |
| JC145 | 630Δ*erm*, *htrA*::CT, P*htrA*-*htrA* | *ermB, catP* | This paper |
| JC148 | 630Δ*erm*, *htrA*::CT, P*htrA*-*htrA^S217A^* | *ermB, catP* | This paper |
| JC166 | 630Δ*erm*, *htrA*::CT, P*htrA-sLuc^opt^* | *ermB, catP* | This paper |
| JC180 | 630Δ*erm htrA^S217A^*, P*htrA-sLuc^opt^* | *catP* | This paper |
| JC204 | 630Δ*erm*, P*cd2243-sLuc^opt^* | *catP* | This paper |
| JC206 | 630Δ*erm*, P*cd2241-sLuc^opt^* | *catP* | This paper |
| JC208 | 630Δ*erm*, P*cd2238-sLuc^opt^* | *catP* | This paper |
| JC212 | 630Δ*erm* *htrA^S217A^*, *Pcd2243-sLuc^opt^* | *catP* | This paper |
| JC214 | 630Δ*erm* *htrA^S217A^*, *Pcd2241-sLuc^opt^* | *catP* | This paper |
| JC216 | 630Δ*erm* *htrA^S217A^*, *Pcd2238-sLuc^opt^* | *catP* | This paper |
| JC218 | 630Δ*erm*, *htrA*::CT*, Pcd2243-sLuc^opt^* | *catP* | This paper |
| JC224 | 630Δ*erm*, *htrA*::CT*, Pcd2241-sLuc^opt^* | *catP* | This paper |
| JC228 | 630Δ*erm*, *htrA*::CT P*cd2238*-*sLuc^opt^* | *ermB, catP* | This paper |
| JC261 | 630Δ*erm*, *htrA*::CT, P*htrA*-*htrAΔPDZ* | e*rmB, catP* | This paper |
| JC285 | 630Δ*erm*, P*tet*-*htrA*-*hiBiT^opt^* | *catP* | This paper |
| JC289 | 630Δ*erm*, *htrA*::CT, P*htrA*-*htrA^S217A^ΔPDZ* | *ermB, catP* | This paper |
| MvL008 | 630Δ*erm*, P*htrA-sLuc^opt^* | *catP* | This paper |

^a^ CT; ClosTron insertion [2], encoding *ermB*.

Table S3: Oligonucleotides used in this study

| Name | Sequence (5’→3’) |
| --- | --- |
| CD3284F2 | AAAACATATGaaggacaatttagtaagtaaatctacagg |
| CD3284R2 | aaaaCTCGAGttaTACTTTTTCATATTTTCCATTTTTGATTAC |
| CD3284F5 | AAAACCATGGGCCATCATCATCATCATCATCATCATCATCACAGCAGCGGCGCCACGCCATCTGTTGTAGGTATTACAACAACTAGTGTTGATACTAG |
| CD3284R3 | aaaactcgagttagaaattcacatttattgttactgcttttc |
| CD3284-S217AF | GCAAGTATAAATGCAGGAAATGCTGGAGGACCATTGTTAAATCAA |
| CD3284-S217AR | TTGATTTAACAATGGTCCTCCAGCATTTCCTGCATTTATACTTGC |
| cd3284_N-terminus_fwd | cgtagaaatacggtgttttttgttaccctaGCTTGTGGAATAAGTGGATATTTATTG |
| cd3284_N-terminus_reV | taacaatggtcctccAGCATTTCCTGCATTTATAC |
| cd3284_C-terminus_fwd | aatgcaggaaatgctGGAGGACCATTGTTAAATC |
| cd3284_C-terminus_rev | gggattttggtcatgagattatcaaaaaggGCTTCACATTTTGAACATC |
| cd3284dowR | AACTCACACTTATCTTGGTCTAAGC |
| CD-pHtrAF2 | AAAAGGTACCACTTAAATAATTTTAAATAGTTAAGGTTTAATTAAG |
| CD-pHtrAR2 | AAAAGAGCTCATCTTAATATAATATAAATTATCAAAATATATTTTTCCC |
| oDB0068 | TAGGATCCGGTTAGAAATTCACATTTATTGTT |
| cd3284upsF2 | GTTCAGAGGAAGTAGACTTAGATGG |
| cd3284upsF3 | CTATGGGTCAATACATACAAGATGG |
| CDPcd2243F | AAAAGGTACCTAAAGCATAAAAATAAGTATTCCTTTCTATC |
| CDPcd2243R | AAAAGAGCTCAAGTTTTACTATTGTAAATTATCAAATATAAC |
| CDPcd2241F | AAAAGGTACCGAAAAATTATAATGGATTTAGAATAATATTG |
| CDPcd2241R | AAAAGAGCTCGTTTCTGAAATTTATTTTCTTGATTTATG |
| CDPcd2238F | AAAAGGTACCTAATATTTAAATGATAAATTGTCTTATAG |
| CDPcd2238R | AAAAGAGCTCTATATAATTATAAAGTCTATTTTTATAC |
| CD3284R2 | aaaaCTCGAGttaTACTTTTTCATATTTTCCATTTTTGATTAC |
| cd3284R6 | aaaactcgagagaaattcacatttattgttactgcttttc |
| cd3284F6 | AAAACATATGATTTATCAAGCAGTTGCAGAAAAAGCTACGCC |
| oDB0067 | CTGAGCTCCTGCAGTAAAGGAGAAAATTTTATGTCAAGAAGAAAGAAAGG |
| oDB0068 | TAGGATCCGGTTAGAAATTCACATTTATTGTT |
| oWKS1070 | GTCTTGGATGGTTGATGAGTAC |
| oWKS1071 | TTCCTAATTTAGCAGCAGCTTC |
| oWKS1387 | CAGATGAGGGCAAGCGGATG |
| oWKS1388 | CGTCGGTGAGCCAGAGTTTC |
| oWKS1537 | TAGGGTAACAAAAAACACCG |
| oWKS1538 | CCTTTTTGATAATCTCATGACC |
| cd3284ITF | CAAAAAACAGTTACTCAAGGGATAATAAG |
| cd3284ITR | CTTATTATCCCTTGAGTAACTGTTTTTTG |
| oWKS1539 | GGATTTCACATTTGCCGTTTTGTAAAC |

Table S4: Plasmids used in this study

| Name | backbone | insert | resistance | purpose | Reference |
| --- | --- | --- | --- | --- | --- |
| pJC014 | pET16B | *htrA*ΔN30 | Amp | Inducible expression in *E. coli* | pVW001 [1] |
| pJC051 | pET16B | *htrA*ΔN30^S217A^ | Amp | Inducible expression in *E. coli* | pJV001 [1] |
| pJC057 | pET16B | *htrA*ΔN64 | Amp | Inducible expression in *E. coli* | This paper |
| pJC058 | pET16B | *htrA*ΔN64^S217A^ | Amp | Inducible expression in *E. coli* | This paper |
| pJC076 | pMTLsc7315 | *htrA*^S217A^ | Cam | ACE mutagenesis to create chromosomal mutant | This paper |
| pJC080 | pRPF185 [3] | P*htrA*-*htrA* | Cam | Complementation of cd3284 knockout | This paper |
| pJC081 | pRPF185 [3] | P*htrA*-*htrA^S217A^* | Cam | Complementation of cd3284 knockout | This paper |
| pJC085 | pAP24 [4] | P*htrA* | Cam | Nanoluciferase reporter assay | This paper |
| pJC086 | pAP24 | P*cd2243* | Cam | Nanoluciferase reporter assay | This paper |
| pJC087 | pAP24 | Pcd2241 | Cam | Nanoluciferase reporter assay | This paper |
| pJC088 | pAP24 | Pcd2238 | Cam | Nanoluciferase reporter assay | This paper |
| pJC102 | pRPF185 | P*htrA*-*htrA*ΔPDZ | Cam | Complementation of cd3284 knockout | This paper |
| pJC105 | pET16B | *htrA*ΔN30ΔPDZ | Amp | Inducible expression in *E. coli* | This paper |
| pJC109 | pET16B | *htrA*^S217A^ΔN30ΔPDZ | Amp | Inducible expression in *E. coli* | This paper |
| pJC113 | pAF302 [5] | htrA | Cam | HiBiT extracellular Detection system | This paper |
| pJC115 | pRPF185 | P*cd3284*-*cd3284*ΔPDZ S217A | Cam | Complementation of cd3284 knockout | This paper |
| pBC013 | pET16B | *cd3284*ΔN55 | Amp | Inducible expression in *E. coli* | This paper |
| pBC015 | pET16B | *cd3284*ΔN55 S217A | Amp | Inducible expression in *E. coli* | This paper |
